# A Strategy to Assess the Cellular Activity of E3 Ligases against Neo-Substrates using Electrophilic Probes

**DOI:** 10.1101/2020.08.13.249482

**Authors:** Benika J. Pinch, Dennis L. Buckley, Scott Gleim, Scott M. Brittain, Laura Tandeske, Pier Luca D’Alessandro, Edward P. Harvey, Zachary J. Hauseman, Markus Schirle, Elizabeth R. Sprague, William C. Forrester, Dustin Dovala, Lynn M. McGregor, Claudio R. Thoma

**Affiliations:** Novartis Institutes for BioMedical Research, Cambridge, MA 02139; Emeryville, CA 94608, USA

**Author notes:** Correspondence (C.R.T.); (L.M.M.); (D.D.); (W.C.F.).

## Abstract

Targeted protein degradation is a rapidly developing therapeutic modality that promises lower dosing and enhanced selectivity as compared to traditional occupancy-driven inhibitors, and the potential to modulate historically intractable targets. While the well-characterized E3 ligases CRBN and VHL have been successfully redirected to degrade numerous proteins, there are approximately 600 predicted additional E3 family members that may offer improved activity, substrate selectivity, and/or tissue distribution; however, characterizing the potential applications of these many ligases for targeted protein degradation has proven challenging. Here, we report the development of an approach to evaluate the ability of recombinant E3 ligase components to support neo-substrate degradation. Bypassing the need for hit finding to identify specific E3 ligase binders, this approach makes use of simple chemistry for Covalent Functionalization Followed by E3 Electroporation into live cells (COFFEE). We demonstrate this method by electroporating recombinant VHL, covalently functionalized with JQ1 or dasatinib, to induce degradation of BRD4 or kinase targets, respectively. Furthermore, by applying COFFEE to SPSB2, a SOCS box and SPRY-domain E3 ligase that has not previously been redirected for targeted protein degradation, we validate this method as a powerful approach to define the activity of previously uncharacterized ubiquitin ligases against neo-substrates.

## INTRODUCTION

Targeted protein degradation (TPD) involves co-opting the cell’s endogenous protein turnover machinery by redirecting a ubiquitin E3 ligase to engage a protein of interest (POI) and promote its subsequent proteolysis at the proteasome. This emerging therapeutic modality promises to expand the landscape of druggable targets by enabling the selective removal of proteins that have been historically difficult to modulate with small molecules, such as proteins with scaffolding functions, or those in closely related families. For example, degradation of the BAF complex component, BRD9, revealed an essential scaffolding role in leukemia and synovial sarcoma that was not observed with small molecule bromodomain inhibitors (Brien et al., 2018; Remillard et al., 2017); and a dual CDK4/6 inhibitor was transformed into selective CDK4 or CDK6 degraders by linking the non-selective inhibitor, palbociclib, to the cereblon (CRBN)-recruiting ligand, thalidomide (Jiang et al., 2019). Furthermore, degraders offer sub-stoichiometric catalytic activity, enabling efficacy at lower doses than their parental inhibitors (Li et al., 2020).

While there are approximately 600 ubiquitin E3 ligases spanning many structural families, ligands for just two substrate receptors, CRBN and von Hippel-Lindau (VHL), have driven rapid expansion of this novel pharmacology, while also limiting its broader applications to other E3s. (Schapira et. al. 2019). Recent work highlights the untapped potential of the E3 family for TPD, with reports that previously unexplored E3 ligase components can be hijacked to induce neo-substrate degradation, such as RNF114 (Spradlin et al., 2019), DCAF16 (Zhang et al., 2019), RNF4 (Ward et al., 2019), and DDB1 (Slabicki et al., 2020). Expanding the TPD toolkit to include ligands for additional E3 ligases could enable tissue- and disease-specific degradation according to E3 ligase expression patterns (Schapira et al., 2019), could broaden the landscape of degradable targets by presenting distinct E3 topologies for ternary complex formation, and/or could offer E3 ligands with improved physicochemical properties.

Structural studies and molecular dynamics simulations have provided insight regarding the dynamics of Cullin-RING E3 ligase-mediated ubiquitination of substrates docked at the receptor. Movement of the flexible RBX/ubiquitin E2 arm and E3 substrate binding domain facilitates scanning of the substrate’s surface accessible lysines for ubiquitin addition, and suggests that the geometry of the substrate-E3 interaction is a critical determinant of efficient ubiquitin transfer (Liu et al., 2009). To our knowledge, there are no reported assays that test the suitability of specific sites on E3 ligases for pharmacological redirection to degrade neo-substrates in living cells. Evaluating the utility of an E3 ligase for TPD requires either genetic engineering of the E3 ligase of interest (Ottis et al., 2017), or the discovery of an E3 ligand followed by the incorporation of that ligand into a bifunctional degrader through additional rounds of medicinal chemistry. We therefore sought to develop a TPD assay to test the “hijackability” of an E3 ligase prior to resource-intensive genetic manipulation or hit-finding chemistry campaigns.

Over 400 E3 ligases have at least one cysteine, making them potentially amenable to covalent targeting. Accordingly, several covalent E3 ligase ligands have been reported and successfully incorporated into bifunctional degraders (Ward et al., 2019; Spradlin et al., 2019; Zhang et al., 2019). While identifying covalent ligands for E3 ligases has proven relatively straightforward, non-optimized covalent ligands often lack selectivity, resulting in off-target activity. While this can be overcome by selective covalent labeling of the purified, recombinant E3, the resulting functionalized E3 is not expected to be cell permeable. In order to develop a probe that would engage various structurally unrelated E3 ligases, and that would enable selective cellular readouts, we sought to append recombinant E3 ubiquitin ligases with covalent probes, and then electroporate the labeled proteins into cells for TPD exploratory studies. Previous reports showcase the utility of electroporation as a relatively gentle method for introducing nucleotides (Tsong et al., 1989), peptides (Schönenberger et al., 2011), proteins (Alex et al., 2019), antibodies (Clift et al., 2018), and various other cell-impenetrant substances into live cells.

In this work, we demonstrate that E3 ligases, including VHL and SPSB2, can be functionalized via their solvent-exposed cysteines using a simple maleimide warhead linked to either the BRD4 ligand, JQ1, or the multi-kinase inhibitor, dasatinib. We further show that the resulting functionalized recombinant E3 ligases can then be electroporated into live cells to form functional E3 ubiquitin ligase complexes capable of catalyzing POI degradation. This method, Covalent Functionalization Followed by E3 Electroporation (COFFEE), provides proof-of-concept information about ligase hijackability that can facilitate the prioritization of E3 ligases for TPD studies and ligand-finding campaigns.

## RESULTS

### Electroporation of Recombinant E3 Ligases Yields Functional CRL Complexes

To identify E3 ligases potentially amenable to COFFEE, we used a bioinformatics approach to identify all E3 ligase components containing cysteines. We then filtered out E3 ligases that catalyze direct transfer of ubiquitin from an E2 to the target protein via an active site cysteine, such as HECT E3s, since modification of the reactive cysteine would presumably inhibit E3 ligase activity (Chen et al., 2018). Finally, we prioritized E3 ligases based both on the availability of structural data, and on a low total cysteine count in order to minimize multiple labeling events (Fig 1A). Using this approach, VHL emerged as a top candidate for functionalization with neo-substrate ligands, having only two cysteines and a well-characterized three-dimensional structure (Hon et al., 2002). Furthermore, as a validated E3 ligase for TPD applications (Bondeson et al., 2015; Zengerle et al., 2015; Gechijian et al., 2018; Bond et al., 2020), VHL presented an ideal proof-of-concept E3 ligase for COFFEE. We therefore expressed purified recombinant VHL in complex with the Cullin-RING ligase (CRL) adaptor proteins, Elongin B (EloB) and Elongin C (EloC), which are required for protein stability. While native VHL exists as two isoforms due to an alternate translation initiation site, we produced the highly expressed and biologically active short isoform (Iliopoulos et al., 1998). To avoid functionalization of the solvent exposed Cys89 of EloB, we also expressed and purified a serine-substituted version, VHL/EloB^C89S^/EloC.

**Figure 1.**
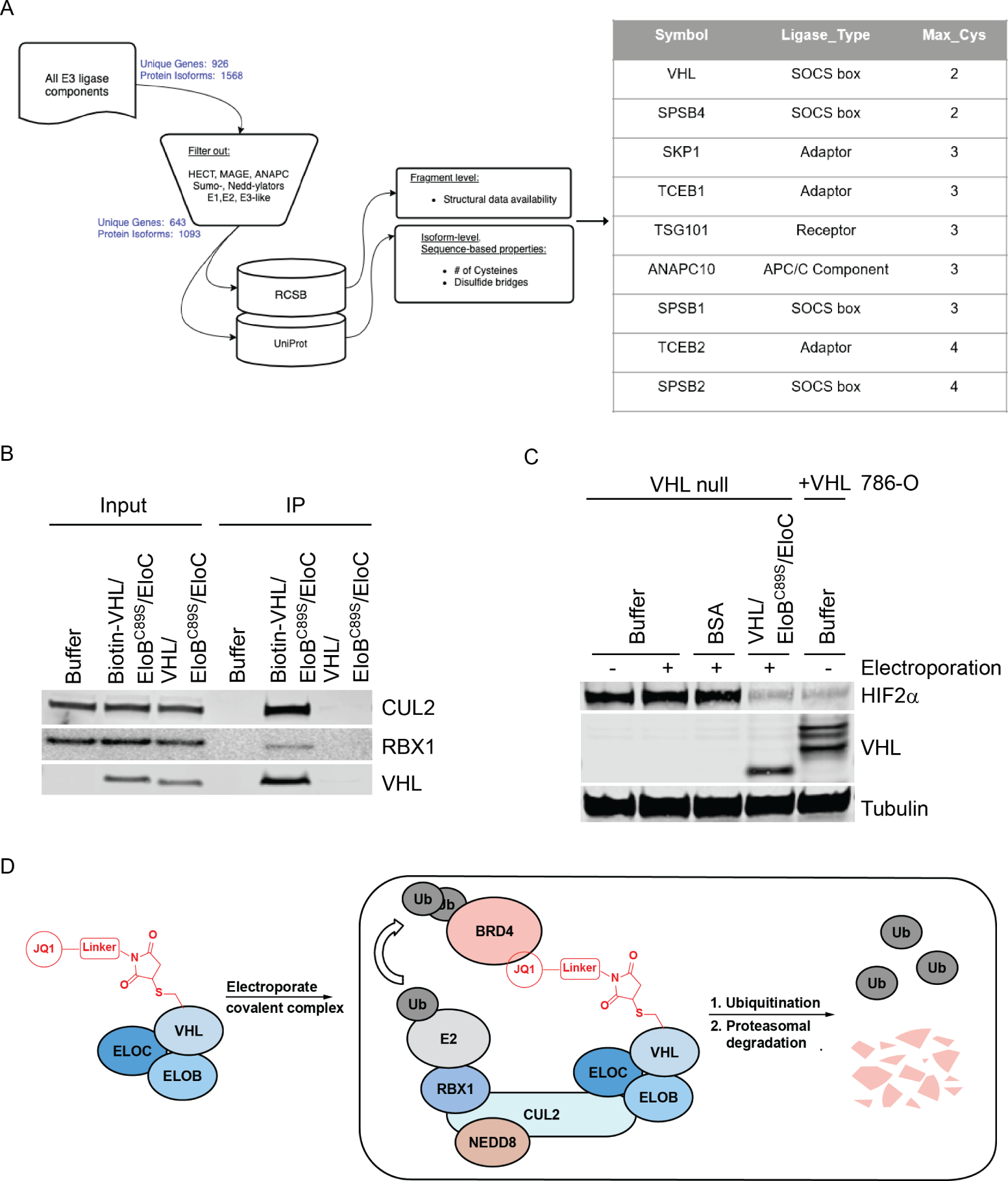
Electroporated VHL is Functional. A. E3 ligase components with potentially ligandable cysteines were identified using a bioinformatics approach. B. Co-IP with streptavidin agarose resin in 786-O cells following electroporation of buffer, biotin-VHL/EloB^C89S^/EloC, or VHL/EloB ^C89S^/EloC. C. Immunoblot analysis following electroporation of buffer, BSA, or VHL/EloB^C89S^/EloC into 786-O cells, as compared to basal HIF2α expression levels in 786-O cells transduced with VHL. D. Schematic depiction of COFFEE workflow, as applied to VHL.

While prior reports demonstrate that electroporated recombinant proteins are functional and exhibit native localization (Alex et al., 2019), to our knowledge, it has not been previously demonstrated that electroporated E3 ligases can associate with endogenous components to form functional ubiquitination complexes. Therefore, following optimization of the electroporation conditions using fluorescently labeled BSA (Fig. S1), we sought to assess whether electroporated VHL/EloB^C89S^/EloC engages Cullin 2 (Cul2)-Rbx1 to form the multimeric CRL2^VHL^ complex. To this end, we performed a co-immunoprecipitation (co-IP) following electroporation of biotin-VHL/EloB^C89S^/EloC (Fig. S1) into the renal cell carcinoma 786-O cell line, which harbors a *VHL* mutation and therefore does not express VHL protein (Iliopoulos et al., 1995). Biotin-VHL/EloB^C89S^/EloC pulled down Cul2 and Rbx1, thereby demonstrating successful assembly of the full CRL2^VHL^ complex following electroporation. By contrast, electroporation of negative controls, including buffer alone or non-biotinylated VHL/EloB^C89S^/EloC, did not pull-down Cul2 or Rbx1 (Fig. 1B).

We next evaluated whether electroporated VHL/EloB^C89S^/EloC resulted in a functional CRL complex, capable of inducing substrate degradation. As compared to non-electroporated cells or cells electroporated with recombinant BSA as a negative control, electroporation of VHL/EloB^C89S^/EloC into 786-O cells decreased levels of VHL’s canonical substrate, HIF2α (Tarade et al., 2019), matching the reduced HIF2α expression observed in 786-O cells expressing both isoforms of VHL from a lentiviral vector (Fig. 1C). Electroporation of WT VHL/EloB/EloC similarly reduced HIF2α expression in 786-O cells (Fig. S2), highlighting that the EloB^C89S^ mutant does not affect functionality of the CRL complex. Successful electroporation of VHL into cells was monitored by immunoblot using an anti-VHL antibody. Taken together, these data confirm that recombinant E3 ligases can be electroporated into cells to form functional E3 ligase complexes capable of mediating ubiquitination and protein degradation through the canonical proteasome-dependent pathway.

### Electroporation of JQ1-functionalized VHL induces BRD4 degradation

Having confirmed that recombinant, electroporated VHL degrades its native substrate, we next tested whether we could use simple cysteine-maleimide chemistry to extend this approach to neo-substrate degradation (Fig. 1D). VHL contains two cysteines, with Cys77 adjacent to the native substrate binding site, and Cys162 buried at the interface between VHL and EloC (Fig. 2A) (Hon et al., 2002). Given that only Cys77 is solvent exposed, VHL should be suitable for facile single labeling with electrophiles, presenting an opportunity to chemically modify the surface of VHL for TPD studies.

**Figure 2.**
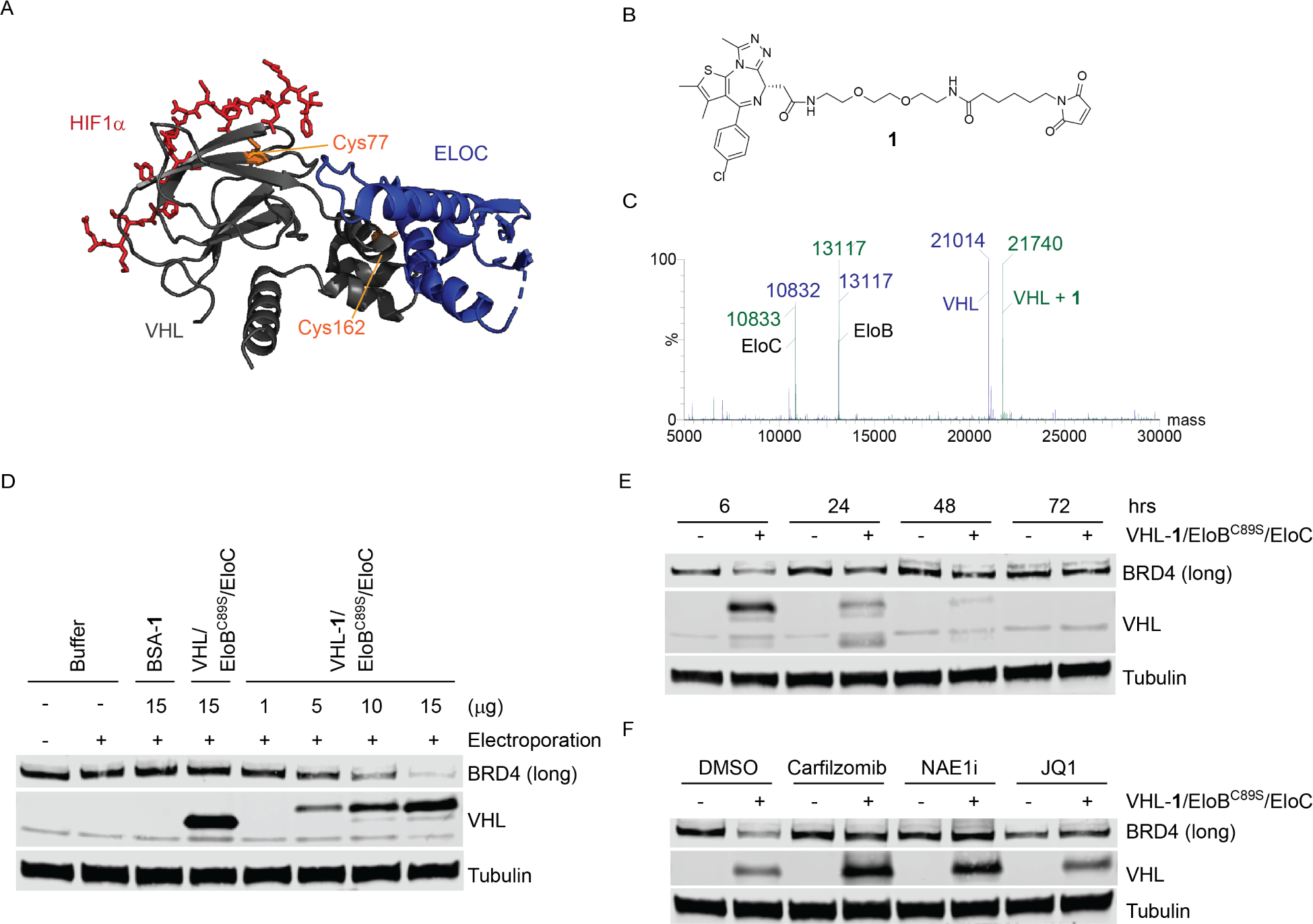
Degradation of BRD4 by JQ1-Functionalized VHL. A. Of the two VHL cysteines (highlighted in orange), Cys77 is adjacent to the native substrate binding site and Cys162 is buried in the interface between VHL and EloC. VHL is shown in gray and Elongin C (ELOC) in blue. Image from PDB 1LQB (Hon et al., 2002). B. Chemical structure of Compound **1**, a maleimide-JQ1 probe. C. Intact mass spectrometry showing complete and specific labeling of VHL by **1**. Unconjugated VHL/EloB^C89S^/EloC is shown in blue, and VHL-**1**/EloB^C89S^/EloC in green. D. Immunoblot analysis showing electroporation of VHL-**1**/EloB^C89S^/EloC into HEK293A cells led to a dose-dependent decrease in BRD4, as compared to cells electroporated with buffer, BSA-**1**, or unlabeled VHL/EloB^C89S^/EloC. Doses are represented as µg of recombinant protein electroporated per well. E. Immunoblot analysis of HEK293A cells pretreated with DMSO, carfilzomib (400 nM), NEDD8 enzyme 1 inhibitor (NAE1i) (1 µM), or JQ1 (10 µM) for 2 hrs, followed by electroporation of VHL-**1**/EloB^C89S^/EloC and co-incubation for 6 hrs. F. Immunoblot analysis of HEK293A cells electroporated with VHL-**1**/EloB^C89S^/EloC, with cells collected at 6, 24, 48, and 72 hrs post-electroporation.

To functionalize VHL Cys77, we synthesized Compound **1**, a bifunctional compound comprised of a cysteine-reactive maleimide connected to the BRD4 ligand, JQ1, via a 2-PEG linker (Fig. 2B) (Filippakopoulous et. al., 2010). Selective and complete single labeling of VHL by **1** was confirmed by intact protein mass spectrometry (Fig. 2C). Electroporation of the resulting VHL-**1**/EloB^C89S^/EloC complex into HEK293A cells led to the dose-dependent loss of BRD4, as assessed by immunoblot analysis (Fig. 2D). Notably, VHL-**1** showed an upward shift in the VHL band by immunoblot as compared to native recombinant VHL, consistent with its increased molecular weight. A panel of negative controls did not affect BRD4 protein levels, including electroporation of buffer alone, unmodified VHL/EloB^C89S^/EloC, or BSA-**1** (Fig. 2D; Fig. S2). Furthermore, incubation of cells with VHL-**1**/EloB^C89S^/EloC without electroporation did not affect BRD4 levels, highlighting that electroporation is necessary for cellular uptake (Fig. S2).

Pretreatment with either the proteasome inhibitor, carfilzomib, or the NEDD8-activating enzyme 1 (NAE1) inhibitor, MLN4924, rescued VHL-**1**/EloB^C89S^/EloC-mediated BRD4 degradation, reflecting a dependence on active CRL activity and proteasomal degradation. Furthermore, pretreatment with an excess of JQ1 (10 µM) also rescued BRD4 degradation, confirming a requirement for BRD4 engagement (Fig. 2F). To evaluate the kinetics of degradation, we performed a time course in HEK293A cells, in which cells were collected 6, 24, 48, or 72 hrs post-electroporation of VHL-**1**/EloB^C89S^/EloC (Fig. 2E). The quantity of electroporated VHL-**1** declined quickly, and was undetectable by 72h, in accordance with VHL’s reported half-life (Zecha et al., 2018; Pozzebon et al., 2013). As expected, the levels of BRD4 recovered as the levels of electroporated VHL-**1** decreased. This highlights that COFFEE must be applied within the half-life of the electroporated protein in order to detect neo-substrate degradation. Taken together, these results confirm that electroporated recombinant VHL-**1** induces the proteasomal degradation of BRD4.

### Electroporation of Dasatinib-functionalized VHL induces degradation of kinase targets

Substrate ubiquitination by cognate E3 ligases or via bifunctional degraders requires the formation of transient complexes. Accordingly, the successful ubiquitination of neo-substrates often reflects their ability to form a de novo protein-protein interface with the recruited E3 ligase (Gadd et al., 2017; Nowak et al., 2018). For example, recent studies show that degradation by bifunctional multi-kinase degraders is dependent on stable ternary complex formation, and that degradation selectivity varies according to the recruited E3 ligase (Tong et al., 2020; Bondeson et al., 2018; Lai et al., 2016). We were therefore interested in using our maleimide-conjugation strategy to extend COFFEE to additional neo-substrate targets by appending VHL with dasatinib, a promiscuous kinase inhibitor that engages 38 kinases with an apparent Ki <100 nM, in addition to its primary target, BCR-ABL (Klaeger et al., 2017; Smith, et. al., 2019).

In order to extend our approach to additional neo-substrates, we synthesized a maleimide-linked dasatinib probe, Compound **2**, using the reported exit vector for dasatinib-based bifunctional degraders (Tong et al., 2020). Compound **2** gave complete single labeling of VHL/EloB^C89S^/EloC, as monitored by intact mass spectrometry (Fig. 3A, 3B). We next evaluated the effects of electroporated VHL-**2**/EloB^C89S^/EloC on ABL1 and Lyn kinase, which are among dasatinib’s top 10 high-affinity targets (Montenegro et al., 2020; Klaeger et al., 2017; Leonard et al., 2016). Electroporation of VHL-**2**/EloB^C89S^/EloC induced the concentration-dependent loss of both ABL1 and Lyn, while electroporation of buffer alone, BSA-**2**, or unmodified VHL/EloB^C89S^/EloC did not affect kinase levels (Fig. 3C). VHL-**2**/EloB^C89S^/EloC-mediated degradation of both ABL1 and Lyn was rescued upon pretreatment with carfilzomib or NAE1i, confirming that degradation occurs through the expected mechanism (Fig. 3D). Degradation of ABL1 was also rescued upon pretreatment with an excess of dasatinib (1 µM), validating a dependence on ABL1 target engagement. Dasatinib treatment alone led to loss of Lyn kinase, likely due to transcriptional effects from broad kinase inhibition at this concentration, or ligand induced protein destabilization. These data highlight that COFFEE can be extended to include ligands against diverse neo-substrates, and can be used to further evaluate the landscape of degradable kinases.

**Figure 3.**
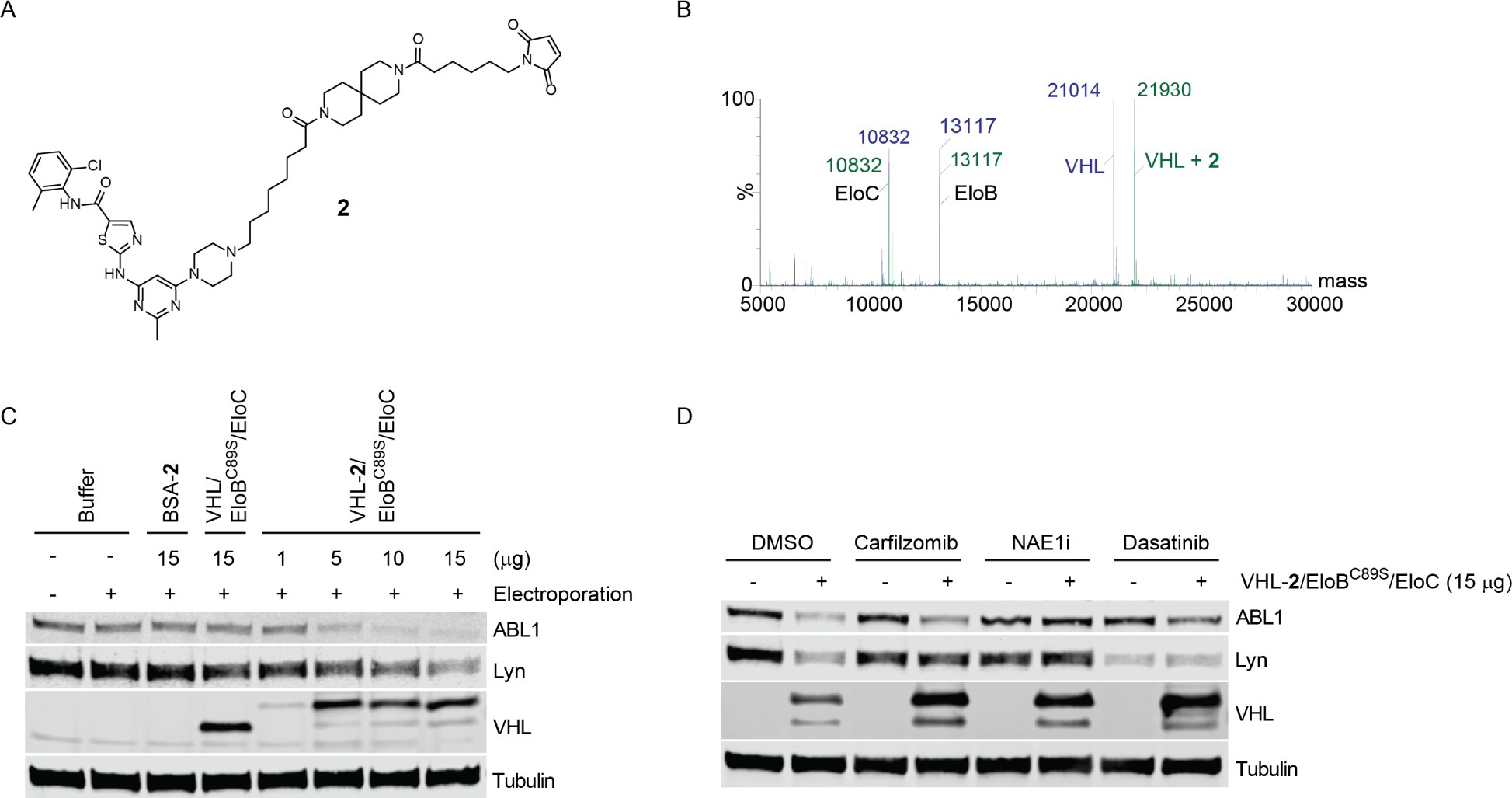
Degradation of Kinases by Dasatinib-Functionalized VHL. A. Chemical structure of Compound **2**, a maleimide-dasatinib probe. B. Intact mass spectrometry confirms complete and specific labeling of VHL by **2**. Unconjugated VHL/EloB^C89S^/EloC is shown in blue, and VHL-**2**/EloB^C89S^/EloC in green. C. Immunoblot analysis showing electroporation of VHL-**2**/EloB^C89S^/EloC into HEK293A cells led to dose-dependent decreases in ABL1 and Lyn, as compared to cells electroporated with buffer, BSA-**2**, or unlabeled VHL/EloB^C89S^/EloC. Doses are represented as µg of recombinant protein electroporated per well. D. Immunoblot analysis of HEK293A cells pretreated with DMSO, carfilzomib (400 nM), NEDD8 enzyme 1 inhibitor (NAE1i) (1 µM), or Dasatinib (1 µM) for 2 hrs, followed by electroporation of VHL**-2/**EloB^C89S^/EloC and co-incubation for 6 hrs.

### SPSB2 is a previously uncharacterized E3 ligase for TPD applications

While VHL provided an ideal proof-of-concept E3 ligase for COFFEE, we next asked whether COFFEE could be applied to a different E3. We therefore expressed and purified another SOCS box E3 ligase containing surface-exposed cysteines, SPSB2, in complex with EloBC (SPSB2/EloB/EloC). SPSB2 is a validated E3 ligase and negative regulator of inducible nitric oxide synthase (iNOS), but has not previously been evaluated for activity against neo-substrates (Kuang et al., 2010). While SPSB2 contains four cysteines, only three are solvent exposed (PDB 5XN3) (You et al., 2017). Whereas stoichiometric reaction with **2** led to multiple labeling events, we established reaction conditions using sub-stoichiometric **2** that led to ∼50% single labeling of SPSB2 Cys53, as monitored by intact protein mass spectrometry (Figs. 4A, 4B, S3). Electroporation of the resulting SPSB2-**2**/EloB/EloC complex into HEK293A cells led to the dose-dependent decrease of ABL1 and Lyn kinases, while electroporation of buffer, BSA-**2**, or unmodified SPSB2/EloB/EloC did not alter kinase levels (Fig. 4C). As expected, SPSB2-**2**/EloB/EloC-mediated kinase degradation was rescued upon pretreatment with carfilzomib or NAE1i (Fig. 4D).

**Figure 4.**
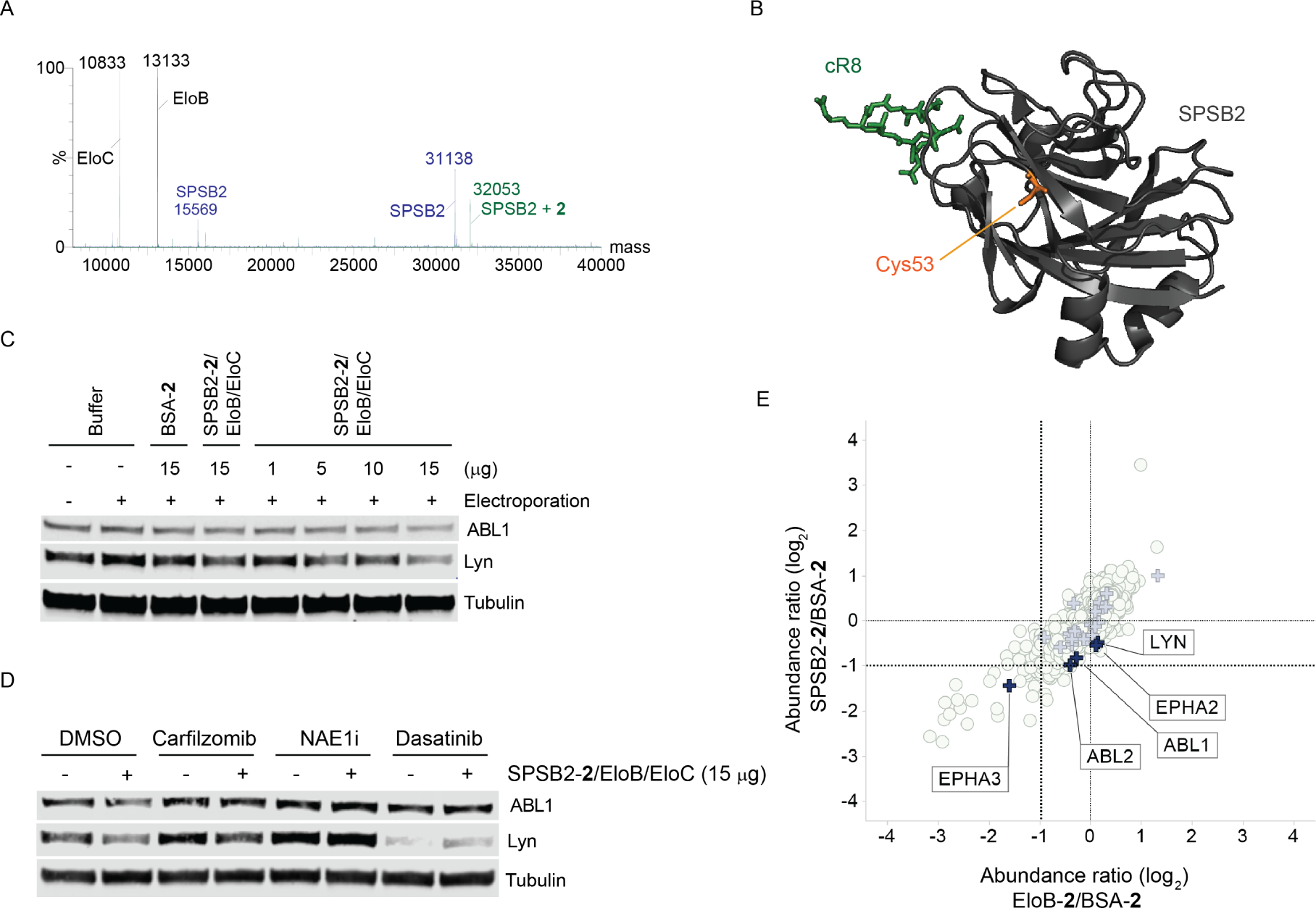
Degradation of Kinases by Dasatinib-Functionalized SPSB2. A. Intact mass spectrometry demonstrates labeling of SPSB2 by **2 (∼50%)**. Unconjugated SPSB2/EloB/EloC is shown in blue, and SPSB2-**2**/EloB/EloC in green. B. SPSB2 Cys53 (highlighted in orange) is solvent exposed and is located near the native substrate binding site. SPSB2 is shown in gray, bound to a reported cyclic peptide inhibitor (cR8), shown in green. Image from PDB 5XN3 (You et al., 2017) C. Immunoblot analysis showing that electroporation of SPSB2-**2**/EloB/EloC into HEK293A cells led to a dose-dependent decrease in ABL and Lyn, while electroporation of buffer, BSA-**2**, or SPSB2/EloB/EloC did not affect kinase abundance. Doses are represented as µg of recombinant protein electroporated per well. D. Immunoblot analysis of HEK293A cells pretreated with DMSO, carfilzomib (400 nM), NEDD8 enzyme 1 inhibitor (NAE1i) (1 µM), or Dasatinib (1 µM) for 2 hrs, followed by electroporation of SPSB2**-2/**EloB/EloC and co-incubation for 6 hrs. E. Quantitative mass spectrometry proteomics reveals that electroporated SPSB2-**2**/EloB/EloC promotes the degradation of dasatinib targets, including EPHB2, EPHB4, ABL1, and ABL2. EloB-**2** weakly promotes the degradation of EPHB2 and EPHB4, suggesting the potential of this E3 adaptor protein to promote the targeted degradation of neo-substrates. Identified targets of dasatinib (Ki < 10 nM) are depicted as blue crosses.

Next, we performed a quantitative proteomics experiment to ask whether other targets of dasatinib might also be degraded by SPSB2-**2**/EloB/EloC. For this experiment, we also selectively modified EloB Cys89 with **2** in order to evaluate the ability of this adaptor component to direct neo-substrate degradation. To do so, we expressed a complex of VHL^C77S^/EloB/EloC, in which EloB Cys89 is the only reactive cysteine (Fig. S3). HEK293A cells were electroporated with SPSB2**-2**/EloB/EloC, VHL^C77S^/EloB-**2**/EloC, or BSA-**2** and incubated for six hours. The resulting lysates were isotopically labeled with TMT reagents and analyzed by nano-LC/MS/MS. The protein abundance ratios (log2) were calculated for SPSB2-**2** vs. BSA-**2** and for EloB-**2** vs. BSA-**2**. The comparison of these ratios demonstrates that electroporation of SPSB2-**2** led to degradation of EPHB2, EPHB4, ABL1, and ABL2, all known targets of dasatinib, as well as moderate degradation of Lyn kinase, in agreement with the immunoblot results (Fig. 4D). Electroporation of VHL^C77S^/EloB-**2**/EloC also led to a modest decrease in the abundance of EPHB2 and EPHB4, highlighting the potential for this E3 adaptor protein to directly mediate neo-substrate degradation. Many other dasatinib targets (Klaeger et al., 2017) were identified in this experiment but were not degraded by SPSB2-**2**/EloB/EloC or by VHL^C77S^/EloB-**2**/EloC.

## DISCUSSION

Recent advances in human genetics research using genome-wide CRISPR loss-of-function or RNAi screens have provided insight into protein dependencies across various diseases (McDonald et. al., 2017; So et al., 2019). Pharmacological efforts to modulate targets by a similar loss-of-function approach have fueled attempts to adapt targeted protein degradation as complementary to classic orthosteric inhibition. However, the field of targeted protein degradation must gain a better understanding of the rules that govern productive combinations of E3 ligase components, target proteins, and chemical matter.

The validation of E3 ligases beyond CRBN and VHL opens the door to novel E3 ligand discovery, which could provide an avenue for optimizing bifunctional degraders beyond the standard “linkerology”, and could expand the landscape of degradable targets. Furthermore, while VHL and CRBN are ubiquitously expressed, recruiting E3 ligases with tissue-or disease-specific expression could improve degradation selectivity and minimize compound-associated toxicity. Therefore, a method to evaluate the ability of functionalized recombinant E3 ligases to catalyze degradation of neo-substrates of interest would enable rapid prioritization of E3s for follow-up ligand finding campaigns. Here, we report the development of such a method, termed Covalent Functionalization Followed by E3 Electroporation (COFFEE), to assess the ability of covalently functionalized recombinant E3 ligases to promote neo-substrate degradation. As proof-of-concept, we demonstrated that electroporated VHL-JQ1 or VHL-dasatinib conjugates successfully induced the proteasome-dependent degradation of BRD4 and kinase targets, respectively. We further extended this approach to validate neo-substrate degradation by SPSB2, demonstrating that this E3 ligase can also be hijacked for TPD applications. There are existing crystal structures of SPSB2 bound by cyclic peptide inhibitors, highlighting the chemical tractability of this E3 (You et al., 2017). Furthermore, the SPSB2 cysteine labeled using COFFEE, Cys53, is unique amongst the SPSB family members (Fig. S3), making the development of selective, covalent SPSB2 ligands another possible path to ligand discovery. Finally, we also demonstrated modest neo-substrate degradation by an EloB-dasatinib conjugate, highlighting that this E3 adaptor component can catalyze degradation, potentially bypassing the need for a substrate receptor. Although we studied just three E3 ligase components, COFFEE relies on simple chemistry, which can be applied to any recombinant E3 ligase component that contains reactive cysteines. Furthermore, the development of a broader set of maleimide-linked ligands, of varying linker lengths and polarities, would further extend the capabilities of COFFEE given that neo-substrate degradation is often target-and linker-dependent (Li et al., 2020).

It is likely that many E3 ligase components possess multiple reactive cysteines, and therefore site-specific modification of such ligases may require the generation of “cysteine-scrubbed” mutant proteins. Such a series of mono-cysteine constructs could enable researchers to evaluate the ability of multiple faces of an E3 ligase component to support targeted protein degradation, as well as the orientation-specific selectivity of degradation. We believe the question of optimal orientation of neo-substrate recruitment can be uniquely addressed by COFFEE, as related methods relying on engineered protein complexes do not offer a similar level of resolution. Furthermore, it is conceivable that preliminary exploratory studies using COFFEE could accelerate the subsequent path to bifunctional degrader development by directly validating particular E3 cysteines for the generation of corresponding covalent ligands or, more generally, by pinpointing E3 binding pockets suitable for ligand finding campaigns. In addition, the composition of the maleimide-linked probes used for COFFEE could inform optimal bifunctional degrader linker length to accelerate medicinal chemistry optimization efforts. We envision that future applications of COFFEE to diverse E3 ligase components will facilitate the expansion of the TPD toolkit beyond ligands for the widely characterized CRBN and VHL, and will accelerate our understanding of the requirements for efficient and selective neo-substrate degradation.

## Acknowledgements

The authors would like to thank Lei Xu for kindly providing cell lines, and Seth Carbonneau, Michael Salcius and Jennifer Lipps for advising on protocols. Also, many thanks to Catherine Dubreuil and the Harvard Program in Therapeutic Science for enabling an internship for B.J.P., E.P.H., and Z.J.H.

## Author Contributions

L.M.M. and B.J.P. wrote the manuscript, with edits from the team. B.J.P. performed the biological experiments. D.L.B. and P.L.D. synthesized the maleimide probes. S.G. performed the bioinformatics analysis. D.D. and L.T. generated the proteins. S.M.B. and L.M.M. performed the proteomics experiment. E.P.H., Z.J.H., M.S., and E.R.S. provided intellectual contributions regarding project directions. W.C.F., D.D., L.M.M., and C.R.T. initiated and supervised the project, and provided experimental support. C.R.T. conceived of the idea.

## METHODS

### Protein Production

The VHL and SPSB2 complexes were expressed from a pETite-based poly-cistronic vector. Constructs were codon-optimized for *E. coli* and synthesized by Twist Biosciences.

C41(DE3) cells (Lucigen) were transformed with the expression plasmid using standard techniques. A single colony was used to start an overnight culture in LB + kanamycin media. This culture was used to inoculate 1 L cultures in Terrific Broth, supplemented with 50 mM sodium phosphate pH 7.0 and 50 ug/mL kanamycin. These cultures grew in Fernbach flasks at 37 degrees C while shaking at 225 rpm, until the OD600 reached approximately 0.625, at which point the temperature was reduced to 20 degrees C and 1 mM IPTG (final) was added to each culture. The cells were allowed to grow overnight.

Cells were harvested by centrifugation at 6,000 x g for 20 minutes and resuspended in Lysis buffer (25 mM Tris pH 8.0, 400 mM NaCl, 1 mM TCEP). Cells were lysed by 3 passes through a cell homogenizer at 18,000 psi. Cell lysate was clarified by centrifugation at 160,000 x g for 2 hours. The whole cell lysate, and then the pellet (following centrifugation) were usually observed to be milky white in appearance, due to the presence of large quantities of inclusion bodies containing the E3 ligase and/or elongins. However, despite the presence of these inclusion bodies (which are removed in this step) there was considerable quantities of soluble protein complex.

Clarified lysate was supplemented with 20 mM imidazole (final) and was then flowed through Ni-NTA resin (5 mL bed volume, gravity fed). The resin was pre-equilibrated with 5 CV of Lysis buffer. After binding of the lysate, the resin was washed with 5 CV of Lysis Buffer, followed by 5 CV of Wash buffer (25 mM Tris pH 8.0, 400 mM NaCl, 40 mM imidazole, 1 mM TCEP), followed by elution with 5 CV of Elution buffer (25 mM Tris pH 8.0, 400 mM NaCl, 500 mM imidazole, 1 mM TCEP).

The IMAC eluate was treated with HRV 3C protease. This reaction mixture was placed into a 10,000 MWCO dialysis cassette and dialyzed against Lysis Buffer overnight at 4 °C. Cleavage was confirmed in the morning by ESI-LC/MS.

The protein was then subjected to reverse-IMAC purification by flowing through 3 mL of Ni-NTA resin pre-equilibrated with Lysis buffer. Upon flow-through, an additional 7 mL of Lysis Buffer was flowed over the resin to recover the remainder of the protein. The flow-through was then concentrated with a centri-con 3,000 MWCO concentrator (Millipore) to approximately 2 mL.

The protein was lastly subjected to Size Exclusion Chromatography utilizing a Superdex 75 16/60 column, pre-equilibrated with SEC Buffer. Protein was loaded and run at 1 mL/min. The complex eluted as a single peak, well separated from the void peak. Pure fractions from this peak were combined and concentrated to 10.7 mg/mL (by nanodrop) and aliquoted

### Cell Culture

HEK293A (Invitrogen) and 293FT (Invitrogen, cat # R70007) were cultured in DMEM supplemented with 10% fetal bovine serum (FBS) and 1% penicillin/streptomycin. 786-O cells (ATCC) were cultured in RPMI 1640 supplemented with 10% FBS and 1% penicillin/streptomycin. All cell lines were cultured at 37°C in a humidified chamber.

### Generation of VHL-expressing 786-O cells

Mcherry in a pENTR plasmid (Thermo Scientific, cat # A10462) and pENTR221-VHL (Thermo Scientific, NM_198156) were Gateway LR cloned into the destination vector pLENTI4/V5 DEST (ThermoFisher Scientific, cat # V49810). Each plasmid was co-transfected with ViraPower(tm) Lentiviral Packaging Mix (ThermoFisher Scientific, cat # K497500) into 293FT cells to make virus particles. Virus particles were filtered through a 0.4 micron filters and used to infect 786-0 cells. Selection was performed with 500 µg/ml of Zeocin for two weeks to obtain 786-0 VHL and 786-0 mcherry pooled cell lines.

### Immunoblotting

Whole cell lysates for immunoblotting were prepared by pelleting cells at 4°C (300 g) for 3 minutes. Cell pellets were then washed 1x with PBS and resuspended in RIPA lysis buffer (VWR, cat # 97063-270) supplemented with protease (Thermo Scientific, cat # A32955) and phosphatase (Thermo Scientific, cat # A32957) inhibitor tablets. Lysates were clarified at 13,200 rpm for 15 min at 4°C prior to quantification by Lowry assay (Bio-Rad cat # 5000113 and cat # 5000114). Whole cell lysates were loaded into 4-20% Criterion TGX Precast 18 well gels (Bio-Rad, cat # 5671094) and separated by electrophoreses at 120 V for 1 hr. The gels were transferred to a nitrocellulose membrane using the Trans-Blot Turbo (Bio-Rad) for 7 minutes and then blocked for 1 hr at room temperature in Odyssey blocking buffer (LICOR Biosciences, cat # 927-50000). Membranes were probed with the appropriate primary antibodies (diluted 1:1000) overnight at 4°C in 20% Odyssey blocking buffer in 1x TBST. Membranes were washed three times with 1x TBST (5 minutes per wash), and then incubated with IRDye goat anti-mouse (LICOR, cat # 926-32210) or goat anti-rabbit (LICOR, cat # 926-32211) secondary antibody diluted 1:10,000 in 20% Odyssey blocking buffer for 1 hr at room temperature. After three 5-minute washes with 1x TBST, the immunoblots were visualized using the ODYSSEY Infrared Imaging System (LICOR). Antibodies used were as follows: Lyn (Cell Signaling, cat # 2796), VHL (Cell Signaling, cat # 68547), HIF2α (Cell Signaling, cat # 7096), α-Tubulin (Cell Signaling, cat # 3973), BRD4 (Cell Signaling, cat # 13440), CUL2 (Invitrogen, cat # 51-1800), RBX1 (Cell Signaling, cat # 4397), and ABL1 (Cell Signaling, cat # 2862).

### Electroporation Optimization by FACS analysis

The pre-programmed 24-well Neon Transfection System protocol using the Neon 10 µL tips (Invitrogen, cat # MPK1096) was used to optimize the electroporation conditions for both HEK293A and 786-O cells. Cells were trypsinized and resuspended in media with 10% FBS but without penicillin/streptomycin prior to counting. Cells were then washed 1x with 5 mL PBS before resuspending in Neon resuspension buffer plus Alexa Fluor 488-labeled BSA, in preparation for 24x 10 µL electroporation reactions (using the Neon pre-programmed 24 optimization conditions), with 250,000 cells and 4.5 ug of Alexa Fluor 488-labeled BSA electroporated per 10 µL reaction. Following electroporation, cells were immediately transferred to a 24-well plate containing 500 µL of pre-warmed media. After a 16 h incubation, cells were washed 1x with 100 µL of PBS prior to trypsinizing (80 µL of TrpLE Express per well). Trypsinized cells were then neutralized with 150 µL of PBS + 20% FBS, mixed by pipetting, and transferred to a 96-well V-bottom plate (Corning, cat # 3894) for FACS analysis. Samples were analyzed using a CytoFLEX benchtop flow cytometer using the FITC channel.

### Electroporation for Immunoblot Analysis

Cells were trypsinized and resuspended in media with 10% FBS but without penicillin/streptomycin prior to counting. Cells were then washed 1x with 5 mL PBS. Each cell pellet was then resuspended in Neon resuspension buffer plus the indicated concentration of recombinant protein to give a final volume of 130 µL per condition (this includes excess volume to avoid bubbles). The cells were then electroporated with the Neon transfection system using the Neon 100 µL kit (Invitrogen, cat # MPK10096) and transferred to a 6-well plate containing 2 mL of pre-warmed media. 2 million cells were electroporated per reaction. The Neon electroporation parameters used were:

1. <u>786-O cells</u>: Pulse Voltage: 1150; Pulse Width: 30; Pulse Number: 2.
2. <u>HEK293A cells</u>: Pulse Voltage: 1400; Pulse Width: 20; Pulse Number: 2.

Cells were collected at the indicated time points following electroporation. To collect cells, each well was washed 1x with 1 mL PBS, followed by trypsinization with 250 µL TrpLE Express (gibco, cat # 12605010). The trypsin was then neutralized with 250 µL media. Cells were washed 1x with 500 µL PBS and then lysed with 50 µL RIPA lysis buffer supplemented with protease and phosphatase inhibitors. Samples were normalized and prepped in 4x LDS (Invitrogen, cat # NP0007) + 10% NuPage sample reducing agent (Invitrogen, cat # NP0009) and boiled for 5 min at 95°C. Lysates were probed for the specified proteins by western blotting.

#### Electroporation Rescue Experiments

Cells were plated in 10 cm plates with 5 million cells per plate in 6 mL of media. The day after plating, each plate was incubated with the indicated pre-treatment condition (DMSO, carfilzomib, MLN4924, JQ1, or dasatinib) at the indicated concentration for 2 hrs. Cells were trypsinized and resuspended in media, followed by electroporation, as detailed above. Following electroporation, cells were transferred to a 6-well plate containing 2 mL of pre-warmed media and were immediately re-treated with the corresponding co-treatment condition, followed by a 6 hr incubation. Cells were then collected as detailed above.

### Biotinylation of VHL/EloB^C89S^/EloC in preparation for Co-Immunoprecipitation

Avi-tagged VHL/EloB^C89S^/EloC was buffer exchanged into 25 mM Tris pH 7.5, 50 mM NaCl using 0.5 mL 7,000 molecular weight cutoff Zeba columns (Thermo Scientific, cat # PI89882), in preparation for biotinylation. 837 ug of VHL/EloBC89S/EloC was incubated with 20 µg of GST-tagged biotin ligase (GST-BirA), 1 mM ATP, 5 mM D-biotin, and 1 mM MgCl_2_ overnight at 4°C (with rotation). The reaction was monitored by intact mass spectrometry. The sample was separated over a 3-minute gradient of 5% to 60% acetonitrile +0.04% trifluoroacetic acid (TFA) in water + 0.05% TFA. Mass spectra are acquired over a mass range of 700 m/z to 3000 m/z. The spectra were then deconvoluted over the depicted mass ranges using Maximum Entropy (MaxEnt). Following reaction completion, the biotinylation mixture was then incubated with 100 µL of a 1:1 glutathione resin (Cytiva, cat # 17075601): buffer (25 mM Tris pH 7.5, 50 mM NaCl) slurry for 1 h at room temperature. Following removal of the glutathione beads by centrifuging in a centrifugal spin column (VWR, modified Nylon membrane, 0.45 µm) at 1.5 rcf for 2 min, the protein was buffer exchanged using a 0.5 mL 7,000 molecular weight cutoff Zeba column into 25 mM HEPES, pH 7.4, 150 mM NaCl, 0.5 mM EDTA. Protein concentration was determined by Nanodrop using the Protein A280 method.

### Co-Immunoprecipitation

786-O cells were trypsinized and resuspended in RPMI media with 10% FBS but without penicillin/streptomycin, prior to counting. Cells were then washed 1x with 5 mL PBS. Each cell pellet was then resuspended in Neon resuspension buffer plus either buffer (25 mM HEPES, pH 7.4, 150 mM NaCl, 0.5 mM EDTA), biotin-VHL/EloBC89S/EloC (15 µg per 100 µL electroporation reaction), or VHL/EloBC89S/EloC (15 µg per 100 µL electroporation reaction) to give a final volume of 130 µL per condition (this includes excess volume to avoid bubbles), with n = 2 electroporation samples per condition. The cells were then electroporated with the Neon transfection system using the Neon 100 µL kit (Invitrogen, cat # MPK10096), with the following electroporation program: Pulse Voltage: 1150; Pulse Width: 30; Pulse Number: 2. Following electroporation, cells were transferred to a 6-well plate containing 2 mL of pre-warmed media and were incubated for 6 hours. 1 million cells were electroporated per 100 µL reaction. To collect cells, each well was washed 1x with 1 mL PBS, followed by trypsinization with 250 µL TrpLE Express (gibco, cat # 12605010). The trypsin was then neutralized with 250 µL media, and the n = 2 samples per condition were pooled. Cells were washed 1x with 500 µL PBS and then lysed with 100 µL Pierce IP lysis buffer (Thermo Scientific, cat # 87787), supplemented with protease and phosphatase inhibitors. Samples were clarified and normalized to 160 µL of 0.65 mg/mL per condition. 10 µL of lysate for each condition was reserved and combined with 10 µL of 2x LDS + 5% NuPage sample reducing agent, boiled for 5 min at 95°C, and loaded as the input control (2.6 µg of total lysate per well of the gel). To the remaining 150 µL of lysate, was added 30 µL of a 1:1 streptavidin agarose resin (Thermo Scientific, cat # 20353): Pierce IP buffer slurry. The lysates were incubated with the streptavidin agarose resin for 1 h at 4°C (with rotation). Beads were washed 4x with 500 µL of Pierce IP buffer. For the final wash, beads were rotated in the washing buffer for 3 min at 4°C. Beads were then pelleted by centrifugation (3,000 rpm for 1 min), dried, and boiled for 5 min at 95°C in 30 µL of 2x LDS + 5% NuPage sample reducing agent. Lysates were probed for the specified proteins by western blotting.

### Covalent Labeling and Monitoring by Intact Mass Spectrometry

Recombinant purified proteins (BSA, VHL/EloB/EloC, VHL/EloB^C89S^/EloC, SPSB2/EloB/EloC, VHL^C77S^/EloB/EloC 50 µM) in 25 mM HEPES, pH 7.4, 150 mM NaCl, 0.5 mM EDTA were incubated with maleimide probes (Compound **1** or **2**; 50 µM) at room temperature. BSA was conjugated to Alexa Fluor 488 using Alexa Fluor 488 C_5_ Maleimide (Invitrogen, cat # A10254). After 2 h, the mass of the resulting complexes was measured by intact protein mass spectrometry, using a Waters Acquity UPLC/ESI-TQD with an Acquity UPLC BEH C4 2.1 × 100 mm, 1.7 um column heated to 80C. The sample was separated over a 3-minute gradient of 5% to 60% acetonitrile +0.04% trifluoroacetic acid (TFA) in water + 0.05% TFA. Mass spectra are acquired over a mass range of 700 m/z to 3000 m/z. The spectra were then deconvoluted over the depicted mass ranges using Maximum Entropy (MaxEnt). If the reaction was incomplete, more maleimide probe was added until completion. The protein was then passed through a 0.5 mL 7,000 molecular weight cutoff Zeba column (Thermo Scientific, cat # PI89882), using 25 mM HEPES, pH 7.4, 150 mM NaCl, 0.5 mM EDTA as the equilibrating buffer, in order to remove any unreacted maleimide probe. Protein concentration was determined by Nanodrop using the Protein A280 method.

### Labeling Site Identification by Peptide Mapping

SPSB2-**2**/EloB/EloC (50 µL of a 5 µM solution) was reduced with 10 mM DTT for 15 minutes at room temperature and alkylated with 25 mM iodoacetamide for 30 minutes at room temperature prior to digestion with 75 ng chymotrypsin at 37C for 4h. The sample was acidified with formic acid to 0.1% v/v and was analyzed on a Thermo Orbitrap Q Exactive Mass Spectrometer (Xcalibur 4.1) coupled to an Easy-nLC 1000 high-performance liquid chromatography system (Thermo Fisher Scientific). A Kasil-fritted trapping column (75 µm x 15 mm) packed with 5 um ReproSil-Pur 120 C18-AQ, was used together with a fused silica spraying capillary pulled to a tip diameter of 8-10 um using a P-2000 capillary puller (Sutter Instruments). The capillary tubing (75 µm I.D.) was packed with a 120 mm separation column comprised of 3 um ReproSil-Pur C18 AQ. Samples (10 µL) were injected onto the trapping column using 0.1% formic acid/2% acetonitrile in water at a flow rate of 2.5 µL/min. Trapped peptides were then introduced into the separation column and eluted at 300 nL/min using a mobile phase A: 2% acetonitrile + 0.1% formic acid in water and a mobile phase B: 98% acetonitrile + 0.1% formic acid in water with the following gradient (3-7% B over 30s followed by 7-45% B over 36 min 30s followed by 45-75% B over 30s followed by 75% B for 5 min). Data were acquired on Q Exactive Mass Spectrometer in data-dependent mode with MS2 triggered on the top 12 precursor ions in the MS1 scan range of 300 – 1500 m/z. Mass spectrometry scans were obtained at 70,000 mass resolution at 200 m/z using a target of 1E6 ions and a maximum fill time of 120 ms. MS2 scans were acquired from 200 to 2000 m/z with normalized collision energy of 25 at a resolution of 17,500 at 200 m/z with an AGC target of 5E4 ions, a maximum fill time of 120 ms. and an isolation window of 2.0 m/z. Dynamic exclusion time 15s, repeat count 1. Charge states of unassigned, +1, and <+5 were excluded. Nanospray voltage 4.5 keV, with capillary temperature 320C, and S-lens RF 50 and sheath gas 10.

### Proteomics

Lysate was generated from previously electroporated HEK293A cell pellets, and 100 ug/sample was reduced, alkylated, and digested with trypsin overnight at 37C. Individual samples were labeled with commercially available TMT Reagents (Thermo Fisher) according to the manufacturer’s protocol. TMT labeled samples were then combined and fractionated by high-pH reversed phase chromatography as previously described.(Spradlin, 2019). The resulting fractions were dried by speed vacuum and reconstituted as 7 fractions. These fractions were analyzed by nanoLC-MS/MS with the instrument and LC conditions described above. Data were acquired on Q Exactive Mass Spectrometer in data-dependent mode with MS2 triggered on the top 12 precursor ions in the MS1 scan range of 300 – 1500 m/z. Mass spectrometry scans were obtained at 35,000 mass resolution at 200 m/z using a target of 1E6 ions and a maximum fill time of 120 ms. MS2 scans were acquired from 200 to 2000 m/z at a resolution of 17,500 at 200 m/z, with an AGC target of 5E4 ions, a maximum fill time of 120 ms, a normalized collision energy of 30 and an isolation window of 2.0 m/z with dynamic exclusion time 10s,and repeat count 1. Isotopes and charge states of unassigned, +1 and <+6 were excluded. Nanospray voltage 4.5 keV, with capillary temperature 320C, and S-lens RF 50 and sheath gas 4.

