## Supplementary Information for "A Strategy to Assess the Cellular Activity of E3 Ligases against Neo-Substrates using Electrophilic Probes" (from another version of this preprint: version 1, 2020-08-14)

**Supplementary Table 1 | Neon Electroporation Optimization**

| | | | | HEK293A | | 786-O | |
|---|---|---|---|---|---|---|---|
| Sample | Pulse Voltage | Pulse Width | Pulse Number | Transfection Efficiency (normalized to condition 1) | % total events (normalized to condition 1) | Transfection Efficiency (normalized to condition 1) | % total events (normalized to condition 1) |
| 1 | None | None | None | 1.00 | 100.00% | 1.00 | 100.00% |
| 2 | 1400 | 20 | 1 | 2.82 | 97.55% | 6.89 | 109.63% |
| 3 | 1500 | 20 | 1 | 5.63 | 102.16% | 9.76 | 94.93% |
| 4 | 1600 | 20 | 1 | 6.38 | 110.26% | 10.79 | 61.44% |
| 5 | 1700 | 20 | 1 | 7.59 | 102.70% | 2.32 | 105.18% |
| 6 | 1100 | 30 | 1 | 4.69 | 76.05% | 5.70 | 81.89% |
| 7 | 1200 | 30 | 1 | 5.88 | 78.83% | 1.87 | 89.35% |
| 8 | 1300 | 30 | 1 | 7.13 | 77.29% | 9.04 | 90.80% |
| 9 | 1400 | 30 | 1 | 8.26 | 77.30% | 1.12 | 70.38% |
| 10 | 1000 | 40 | 1 | 4.82 | 79.27% | 6.21 | 72.54% |
| 11 | 1100 | 40 | 1 | 6.27 | 77.23% | 1.68 | 71.60% |
| 12 | 1200 | 40 | 1 | 7.56 | 57.10% | 11.08 | 60.38% |
| 13 | 1100 | 20 | 2 | 5.31 | 77.76% | 5.86 | 69.61% |
| 14 | 1200 | 20 | 2 | 7.12 | 76.12% | 7.42 | 58.62% |
| 15 | 1300 | 20 | 2 | 9.34 | 61.93% | 8.06 | 51.24% |
| 16 | 1400 | 20 | 2 | 11.25 | 56.64% | 2.29 | 72.77% |
| 17 | 850 | 30 | 2 | 3.09 | 73.29% | 3.69 | 60.27% |
| 18 | 950 | 30 | 2 | 5.37 | 56.45% | 5.72 | 56.71% |
| 19 | 1050 | 30 | 2 | 8.70 | 70.59% | 8.62 | 59.73% |
| 20 | 1150 | 30 | 2 | 11.67 | 70.96% | 11.38 | 47.54% |
| 21 | 1300 | 10 | 3 | 4.46 | 57.41% | 5.18 | 50.81% |
| 22 | 1400 | 10 | 3 | 6.20 | 47.17% | 6.43 | 32.73% |
| 23 | 1500 | 10 | 3 | 2.51 | 60.82% | 7.52 | 18.17% |
| 24 | 1600 | 10 | 3 | 8.68 | 68.50% | 2.47 | 60.98% |

HEK283A or 786-O cells were electroporated with Alexa-Fluor 488-conjugated BSA according to the 24-well pre-programmed Neon electroporation parameters listed. Following a 16 h incubation, cells were analyzed by FACS to determine median signal intensity (FITC channel) and percent total events, normalized to Condition 1 (non-electroporated control).

**SUPPLEMENTARY FIGURES**

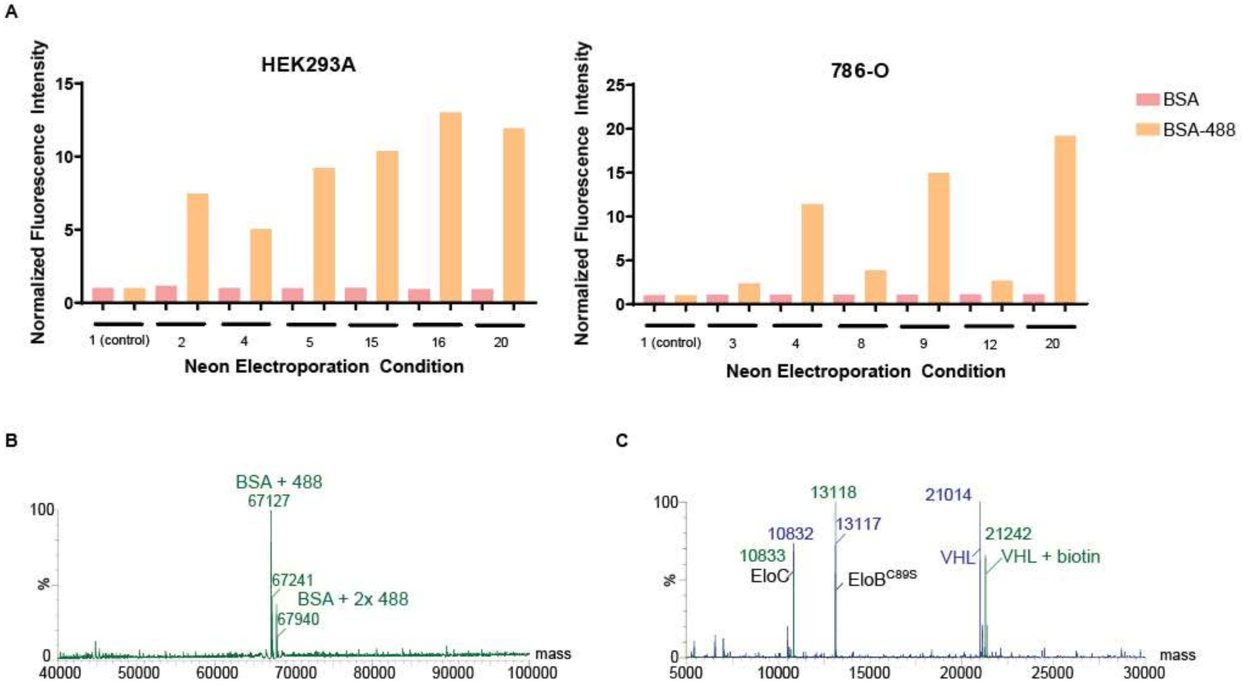


**Supplementary Figure 1 | Electroporation Optimization.** (**A**) Normalized fluorescence intensity in HEK293A (left) or 786-O (right) cells following electroporation with either BSA or Alexa-Fluor 488-conjugated BSA using the indicated Neon pre-programmed optimization electroporation parameters (Supplementary Table 1). Cells were incubated for 16 h prior to FACS analysis. (**B**) Conjugation of AlexaFluor 488 to BSA is confirmed by intact MS. (**C**) Biotinylation of VHL by GST-BirA is confirmed by intact MS, with unbiotinylated VHL/EloB$^{C89S}$/EloC shown in blue, and biotinylated VHL/EloB$^{C89S}$/EloC shown in green.

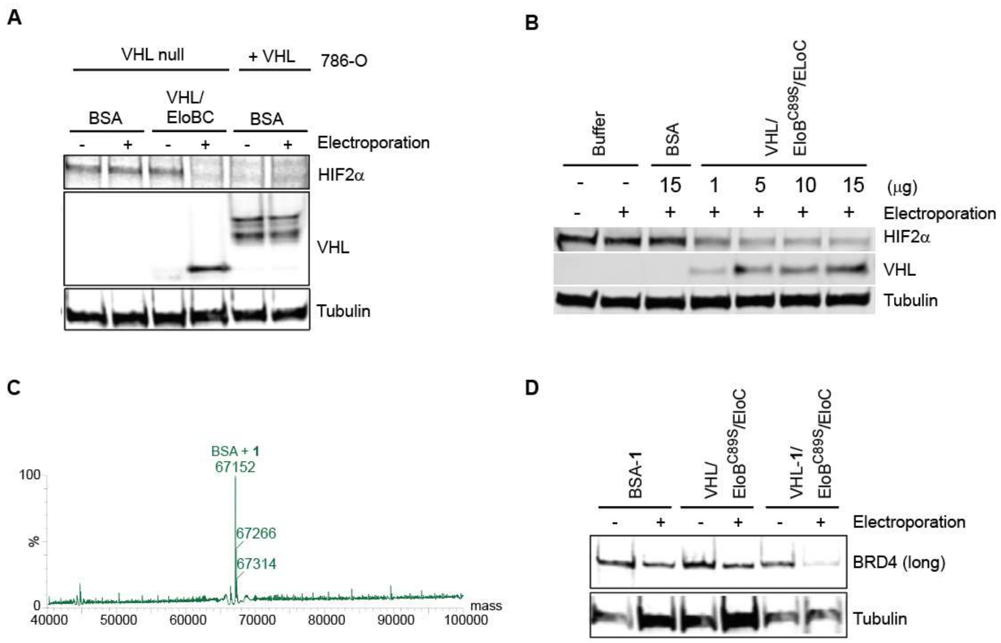


**Supplementary Figure 2 | Electroporated VHL is functional and can be redirected to degrade BRD4 using COFFEE.** (**A**) Immunoblot analysis following electroporation of BSA or WT VHL/EloB/EloC into 786-O cells (VHL null), and electroporation of BSA into 786-O cells transduced with VHL. Cells were collected 15 h post-electroporation. (**B**) Immunoblot analysis following electroporation of BSA or VHL/EloB$^{C89S}$/EloC into 786-O cells at the indicated concentrations (represented as μg of protein electroporated per 100 μL electroporation reaction). Cells were collected 6 h post-electroporation. (**C**) Intact mass analysis confirms labeling of BSA by compound **1**; (**D**) Immunoblot analysis following incubation of BSA-**1**, VHL/EloB$^{C89S}$/EloC, or VHL-**1**/EloB$^{C89S}$/EloC into HEK293A cells – with or without electroporation for each condition. Cells were collected 8 h post-electroporation. 15 μg of protein was electroporated per 100 μL electroporation reaction.

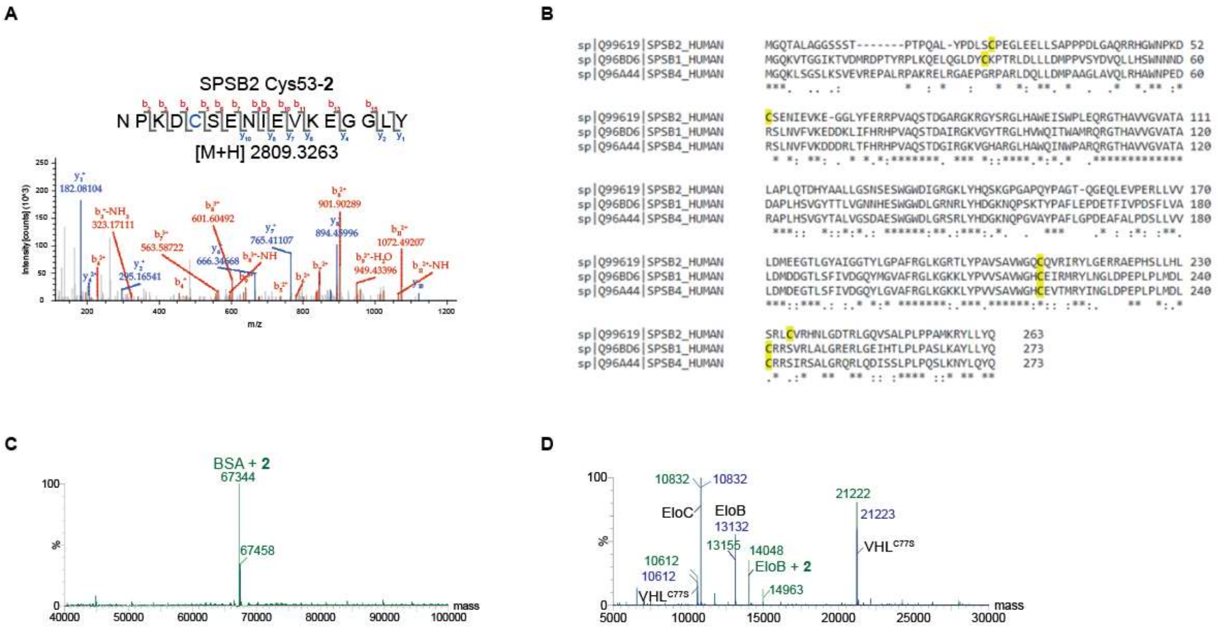


**Supplementary Figure 3 | SPSB2 functionalization with Compound 2.** (**A**) Peptide mapping for SPSB2-**2**/EloBC, showing that compound **2** labels SPSB2 Cys 53. (**B**) Sequence alignment of SPSB family showing that SPSB2 Cys53 is unique amongst the family members. (**C**) Intact mass analysis confirms labeling of BSA by compound **2**. (**D**) Intact mass analysis demonstrates ~50% labeling of EloB by compound **2**, with unlabeled $VHL^{C77S}$/EloB/EloC shown in blue, and labeled $VHL^{C77S}$/EloB-**2**/EloC shown in green.

**SUPPLEMENTARY METHODS: Chemistry Experimental**

**Compound 1 Synthetic Methods:**

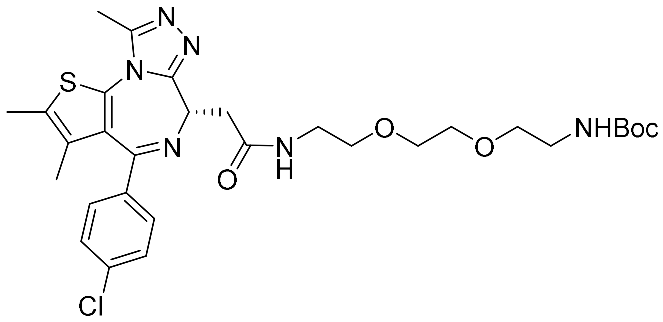


**tert-butyl (S)-(2-(2-(2-(2-(4-(4-chlorophenyl)-2,3,9-trimethyl-6H-thieno[3,2-f][1,2,4]triazolo[4,3-a][1,4]diazepin-6-yl)acetamid)ethoxy)ethoxy)ethyl)carbamate**

(S)-2-(4-(4-chlorophenyl)-2,3,9-trimethyl-6H-thieno[3,2-f][1,2,4]triazolo[4,3-a][1,4]diazepin-6-yl)acetic acid (133.3 mg, 0.333 mmol, 1 eq) and tert-butyl (2-(2-(2-aminoethoxy)ethoxy)ethyl)carbamate (83 mg, 0.333 mmol, 1 eq) were dissolved in DMF (3.3 mL) at room temperature. DIPEA (175 microliters, 1.0 mmol, 3 eq) was added, followed by HATU (127 mg, 0.333 mmol, 1 eq). After 24 hours, the material was diluted with EtOAc and washed with saturated sodium bicarbonate, water and brine. The material was dried over sodium sulfate, filtered and concentrated under reduced pressure. Purification by column chromatography (ISCO, 12 g silica column, 0-15% MeOH/DCM) gave the desired product (235.1 mg, 0.372 mmol, quantitative yield) as a yellow oil.

**$^1$H NMR** (400 MHz, Chloroform-d) δ 7.44 - 7.38 (m, 2H), 7.33 (d, J = 8.7 Hz, 2H), 6.94 (s, 1H), 5.27 (s, 1H), 4.65 (t, J = 6.9 Hz, 1H), 3.63 (s, 4H), 3.62 - 3.47 (m, 7H), 3.42 - 3.28 (m, 3H), 2.67 (s, 3H), 2.40 (s, 3H), 1.71 - 1.62 (m, 3H), 1.45 (s, 9H).

**LCMS**: 631.5 $[M+H]^+$.

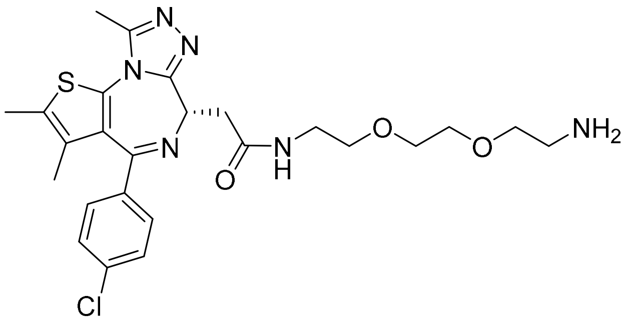


**(S)-N-(2-(2-(2-aminoethoxy)ethoxy)ethyl)-2-(4-(4-chlorophenyl)-2,3,9-trimethyl-6H-thieno[3,2-f][1,2,4]triazolo[4,3-a][1,4]diazepin-6-yl)acetamide hydrochloride salt**

tert-butyl (S)-(2-(2-(2-(2-(4-(4-chlorophenyl)-2,3,9-trimethyl-6H-thieno[3,2-f][1,2,4]triazolo[4,3-a][1,4]diazepin-6-yl)acetamido)ethoxy)ethoxy)ethyl)carbamate (235.1 mg, 0.372 mmol, 1 eq) was dissolved in MeOH (2.5 mL) and 3M HCl (MeOH) (2.5 mL, 7.45 mmol) at room temperature. After 48 hours the material was concentrated under reduced pressure and dried by high vac. The desired product was isolated (166.1 mg, 0.293 mmol, 79%) as a yellow solid and used without further purification.

**$^1$H NMR** (400 MHz, Methanol-d4) δ 7.69 - 7.52 (m, 4H), 5.13 - 5.03 (m, 1H), 3.78 - 3.74 (m, 2H), 3.73 (s, 4H), 3.66 (t, J = 5.6 Hz, 2H), 3.63 - 3.46 (m, 4H), 3.16 (t, J = 5.0 Hz, 2H), 3.04 - 2.91 (m, 3H), 2.54 (s, 3H), 1.75 (s, 3H).

**LCMS**: 533.3 $[M+H]^+$

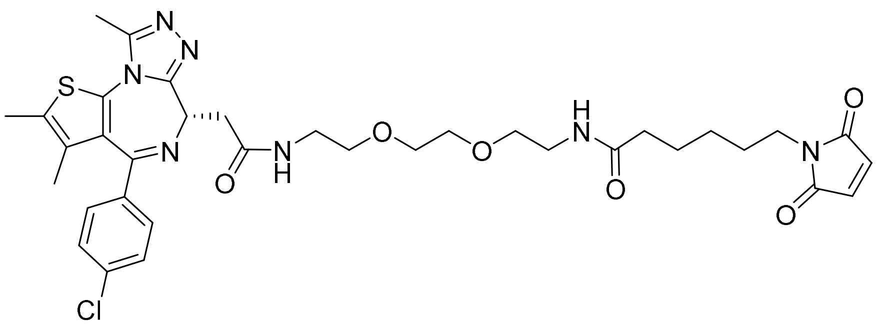

**(S)-N-(2-(2-(2-(2-(4-(4-chlorophenyl)-2,3,9-trimethyl-6H-thieno[3,2-f][1,2,4]triazolo[4,3-a][1,4]diazepin-6-yl)acetamido)ethoxy)ethoxy)ethyl)-6-(2,5-dioxo-2,5-dihydro-1H-pyrrol-1-yl)hexanamide (Compound 1)**

(S)-N-(2-(2-(2-aminoethoxy)ethoxy)ethyl)-2-(4-(4-chlorophenyl)-2,3,9-trimethyl-6H-thieno[3,2-f][1,2,4]triazolo[4,3-a][1,4]diazepin-6-yl)acetamide hydrochloride salt (47.5 mg, 0.84 mmol, 1 eq) and 6-(2,5-dioxo-2,5-dihydro-1H-pyrrol-1-yl)hexanoic acid (17.68 mg, 0.084 mmol, 1 eq) were dissolved in DMF (0.84 mL) at room temperature. DIPEA (44 microliters, 0.251 mmol, 3 eq) was added, followed by HATU (31.8 mg, 0.084 mmol, 1 eq). After 23 hours, the mixture was diluted with EtOAc (50 mL) and washed three times with brine (25 mL). The organic layer was dried over sodium sulfate, filtered and concentrated under reduced pressure. Purification by column chromatography (ISCO, 12 g silica column, 0-15% MeOH/DCM) gave the desired product (41.6 mg, 0.048 mmol, 57% yield) as a yellow oil.

**$^{1}$H NMR** (400 MHz, Chloroform-*d*) δ 7.40 – 7.34 (m, 2H), 7.30 – 7.25 (m, 2H), 7.08 (s, 1H), 6.60 (d, *J* = 12.7 Hz, 3H), 4.65 (dd, *J* = 7.7, 6.3 Hz, 1H), 3.59 (q, *J* = 3.3 Hz, 4H), 3.56 – 3.52 (m, 4H), 3.44 (t, *J* = 5.2 Hz, 2H), 3.41 – 3.34 (m, 4H), 2.66 (s, 3H), 2.38 – 2.32 (m, 3H), 2.10 (t, *J* = 7.6 Hz, 2H), 1.55 (dd, *J* = 9.0, 6.4 Hz, 2H), 1.46 (q, *J* = 7.6 Hz, 2H), 1.20 (dtd, *J* = 7.9, 4.8, 2.1 Hz, 4H).

**LCMS**: 724.5 $[M+H]^{+}$

**Compound 2 Synthetic Methods:**

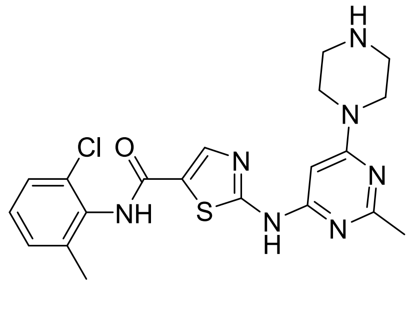

**N-(2-chloro-6-methylphenyl)-2-((2-methyl-6-(piperazin-1-yl)pyrimidin-4-yl)amino)thiazole-5-carboxamide**

A mixture of piperazine (4.37 g, 50.7 mmol, 10.0 eq.), 2-((6-chloro-2-methylpyrimidin-4-yl)amino)-N-(2-chloro-6-methylphenyl)thiazole-5-carboxamide (commercially available from ABCR) (2 g, 5.07 mmol, 1.0 eq.) and DIPEA (1.8 ml, 10.15 mmol, 10.15 eq.) in Dioxane (60 mL) was heated at reflux for 2 hours. After 2 hours the reaction was allowed to reach room temperature and was stirred overnight. Next, the reaction was evaporated and the crude material was triturated in MeOH to afford the desired product (1 g, 2.2 mmol, 44 % yield) as a white solid.

**$^1$H NMR** (400 MHz, DMSO-$d_6$) δ 9.86 (s, 1H), 8.21 (s, 1H), 7.40 (dd, $J$ = 7.5, 2.0 Hz, 1H), 7.36 – 7.21 (m, 2H), 6.02 (s, 1H), 3.44 (t, $J$ = 5.1 Hz, 4H), 3.31 (s, 4H), 2.74 (t, $J$ = 5.0 Hz, 4H), 2.40 (s, 3H), 2.24 (s, 3H).

**LC-MS:** 444.1 $[M+1]^+$

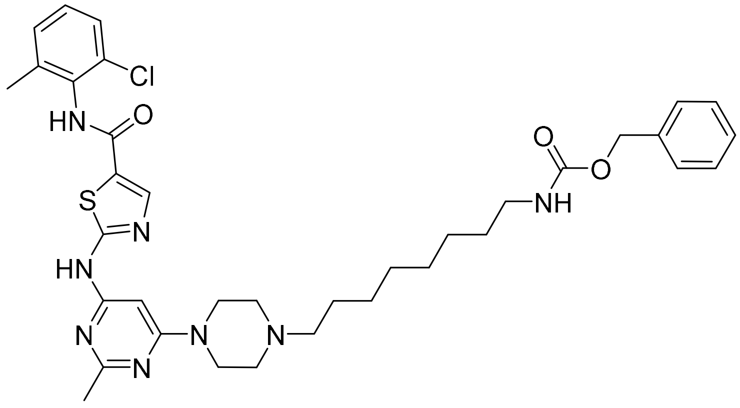


**Benzyl (8-(4-(6-((5-((2-chloro-6-methylphenyl)carbamoyl)thiazol-2-yl)amino)-2-methylpyrimidin-4-yl)piperazin-1-yl)octyl)carbamate**

Acetic acid (0.077 ml, 1.35 mmol, 1.2 eq.) was added to a suspension of N-(2-chloro-6-methylphenyl)-2-((2-methyl-6-(piperazin-1-yl)pyrimidin-4-yl)amino)thiazole-5-carboxamide (500 mg, 1.12 mmol, 1.0 eq.) and benzyl (8-oxooctyl)carbamate (commercially available from Aldlab Chemicals LLC [134477-40-0]) (469 mg, 1.7 mmol, 1.5 eq.) in 50% Methanol/DCM (16.00 mL). After stirring the reaction for 2 hours at room temperature $NaCNBH_3$ (106 mg, 1.7 mmo, 1.5 eq.) was added. The reaction was stirred overnight at room temperature and after evaporated under reduced pressure. The obtained crude material was purified by flash chromatography (silica column 40 g, MeOH/DCM mobile phase to obtain the desired product (550 mg, 0.780 mmol, 69 % yield) as a white solid.

**LC-MS:** 705.4 $[M+1]^+$

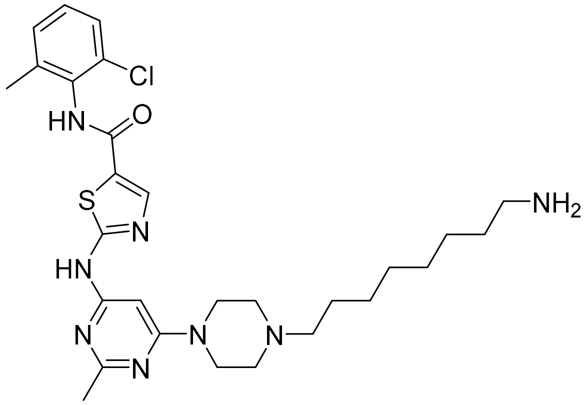


**2-((6-(4-(8-aminooctyl)piperazin-1-yl)-2-methylpyrimidin-4-yl)amino)-N-(2-chloro-6-methylphenyl)thiazole-5-carboxamide**

HCl in MeOH (1.25 M) (2 mL, 2.5 mmol, 8.0 eq.) was added to a solution of benzyl (8-(4-(6-((5-((2-chloro-6-methylphenyl)carbamoyl)thiazol-2-yl)amino)-2-methylpyrimidin-4-yl)piperazin-1-yl)octyl)carbamate (222 mg, 0.31 mmol, 1.0 eq.) in THF (3 mL). The reaction was hydrogenated ($H_2$ 0.1 bar) at working temperature 22-25 °C. After 127 hours, the reaction was filtered through a pad of Hyflo Super Cel. The pad was washed with 50% MeOH/DCM (10 mL) and the combined solution was evaporated to afford the desired product (167 mg, 0.254 mmol, 81 % yield) as HCl salt.

**$^1$H NMR** (600 MHz, DMSO-d6) δ 11.65 (s, 1H), 10.90 (s, 1H), 9.94 (s, 1H), 8.25 (s, 1H), 7.83 (s, 3H), 7.40 (d, J = 7.5 Hz, 1H), 7.32 - 7.22 (m, 2H), 6.15 (s, 1H), 4.33 (s, 2H), 3.55 (d, J = 10.2 Hz, 2H), 3.37 (s, 2H), 3.12 - 2.93 (m, 4H), 2.80 - 2.70 (m, 2H), 2.44 (s, 3H), 2.24 (s, 3H), 1.71 (s, 2H), 1.58 - 1.49 (m, 2H), 1.29 (s, 8H).

**LC-MS:** 571.2 $[M+1]^+$

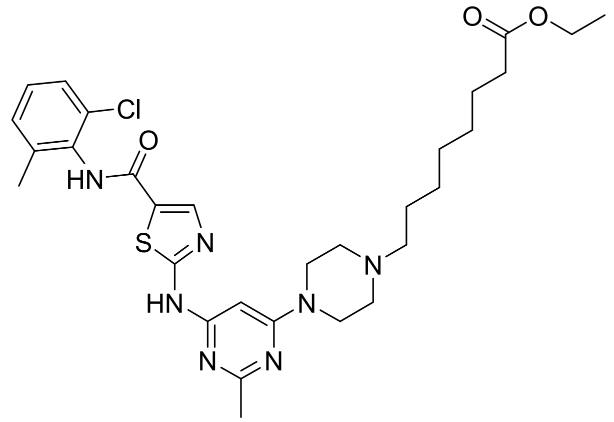


**Ethyl 8-(4-(6-((5-((2-chloro-6-methylphenyl)carbamoyl)thiazol-2-yl)amino)-2-methylpyrimidin-4-yl)piperazin-1-yl)octanoate**

Ethyl 8-bromooctanoate (0.85 g, 3.38 mmol, 1.5 e.q) was added to a mixture of 2-((6-chloro-2-methylpyrimidin-4-yl)amino)-N-(2-chloro-6-methylphenyl)thiazole-5-carboxamide (1 g, 2.2 mmol, 1.0 eq.) and DIPEA (2 mL, 11.45 mmol, 5.2 eq.) in DMF (20 mL). The reaction was heated at 80°C under stirring overnight. Next, the reaction was evaporated under reduced pressure to obtain a residue which was dissolved in Ethyl acetate (200ml) and saturated solution of $NaHCO_3$ (100ml). The organic phase was separated, washed with brine (100 mL x 2) and evaporated to under reduced pressure. The obtained crude material was purified by flash chromatography (silica column 80g, MeOH/DCM mobile phase) to obtain the desired product (1.2 g, 1.9 mmol, 87 % yield) as a white solid.

**LC-MS:** 614.3 $[M+1]^+$

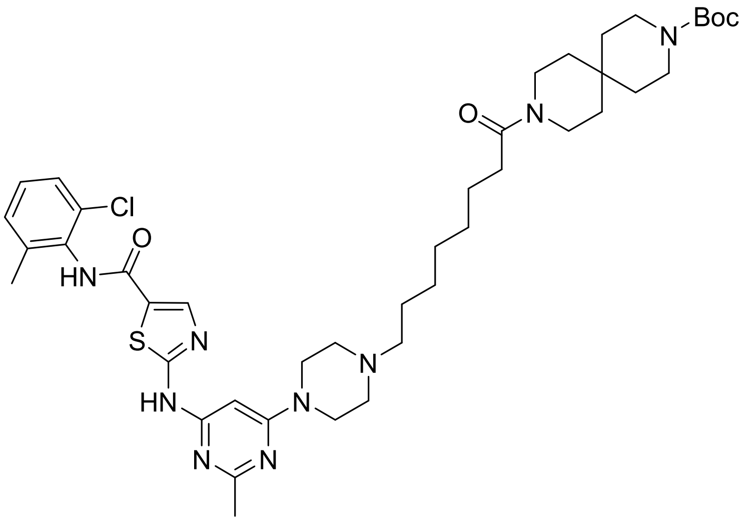

**tert-butyl 9-(8-(4-(6-((5-((2-chloro-6-methylphenyl)carbamoyl)thiazol-2-yl)amino)-2-methylpyrimidin-4-yl)piperazin-1-yl)octanoyl)-3,9-diazaspiro[5.5]undecane-3-carboxylate**

LiOH (2M) (0.68 mL, 1.35 mmol) was added to a suspension of ethyl 8-(4-(6-((5-((2-chloro-6-methylphenyl)carbamoyl)thiazol-2-yl)amino)-2-methylpyrimidin-4-yl)piperazin-1-yl)octanoate (280 mg, 0.45 mmol, 1.0 eq.) in THF (4 mL). After stirring the reaction for 2 days at room temperature, water (1 mL) was added and the obtained solution was lyophilized to afford 8-(4-(6-((5-((2-chloro-6-methylphenyl)carbamoyl)thiazol-2-yl)amino)-2-methylpyrimidin-4-yl)piperazin-1-yl)octanoic acid (302 mg, 0.47 mmol, 100 % yield) as a white solid lithium salt. **LC-MS:** 586.2 $[M+1]^+$

TEA (0.131 mL, 0.9 mmol, 3.0 eq.) and HBTU (356 mg, 0.9 mmol, 3.0 eq.) were added to a solution of the intermediate acid derivative (200 mg, 0.3 mmol, 1.0 eq.) in DMF (2 mL) and the reaction was stirred at room temperature for 30 min. Next, tert-butyl 3,9-diazaspiro[5.5]undecane-3-carboxylate (199 mg, 0.8 mmol, 2.7 eq.) was added. After stirring at room temperature overnight, the mixture was poured in EA (120 mL), washed with a saturated solution of $NaHCO_3$ (80 mL x 3), brine (80mL), dried with $MgSO_4$ and evaporated under reduced pressure. The crude material was purified by flash chromatography (silica column 24 g, MeOH/DCM mobile phase) to obtain the desired product (201 mg, 0.23 mmol, 75 % yield) as a yellow solid.

**LC-MS:** 822.5 $[M+1]^+$

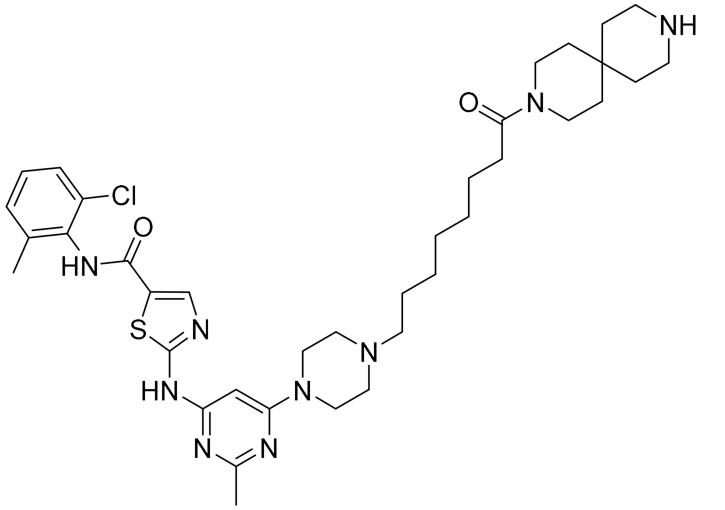

**N- (2-chloro-6-methylphenyl)-2-((2-methyl-6-(4-(8-oxo-8-(3,9-diazaspiro[5.5]undecan-3-yl)octyl)piperazin-1-yl)pyrimidin-4-yl)amino)thiazole-5-carboxamide**

HCl in Dioxane (3M) (5 ml, 15.00 mmol, 30.5 eq.), water (0.250 mL) and MeOH (2 mL) were added to a suspension of tert-butyl 9-(8-(4-(6-((5-((2-chloro-6-methylphenyl)carbamoyl)thiazol-2-yl)amino)-2-methylpyrimidin-4-yl)piperazin-1-yl)octanoyl)-3,9-diazaspiro[5.5]undecane-3-carboxylate (417 mg, 0.49 mmol, 1 eq.) in Dioxane (3 mL). After stirring at room temperature overnight the reaction was evaporated to dryness under reduced pressure. The crude material was purified by reverse phase flash chromatography (C18 50g Gold, ACN/Water mobile phase, TFA modifier), then lyophilized to obtain the desired product (408 mg, 0.425 mmol, 86 % yield) as a white solid TFA salt.

**$^1$H NMR** (400 MHz, DMSO-d6) δ 11.64 (s, 1H), 9.90 (s, 1H), 9.69 (s, 1H), 8.34 (s, 2H), 8.24 (s, 1H), 7.40 (d, J = 7.5 Hz, 1H), 7.33 - 7.22 (m, 2H), 6.15 (s, 1H), 4.36 (d, J = 12.5 Hz, 2H), 3.58 (d, J = 11.7 Hz, 2H), 3.44 - 3.40 (m, 2H), 3.39 - 3.34 (m, 2H), 3.22 (t, J = 12.5 Hz, 2H), 3.11 (s, 2H), 3.02 (s, 4H), 2.49 - 2.48 (m, 2H), 2.45 (s, 3H), 2.28 (t, J = 7.4 Hz, 2H), 2.24 (s, 3H), 1.65 (s, 2H), 1.60 (d, J = 5.1 Hz, 4H), 1.46 (d, J = 16.6 Hz, 4H), 1.38 (t, 2H), 1.34 - 1.25 (m, 6H).

**LC-MS:** 722 $[M+1]^+$

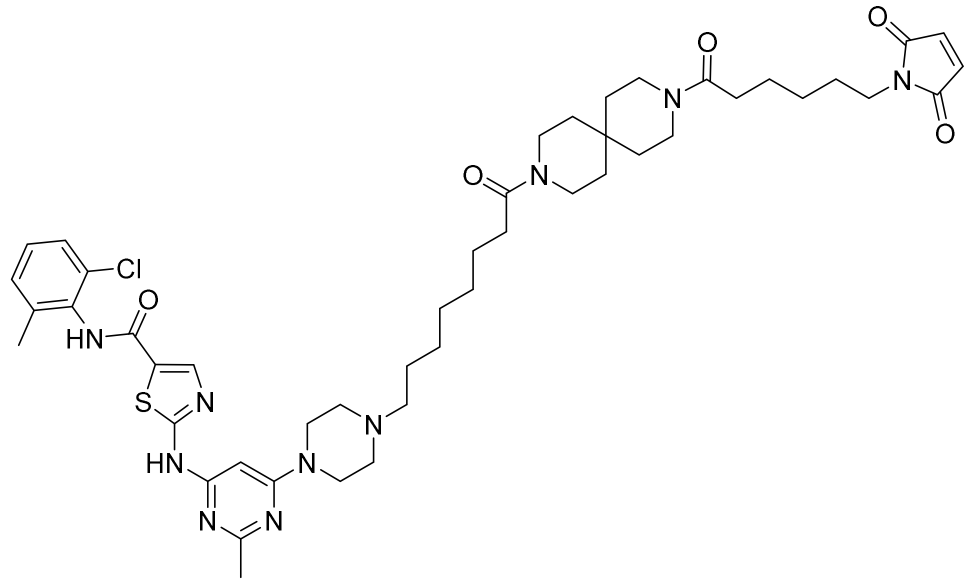


**N-(2-chloro-6-methylphenyl)-2-((6-(4-(8-(9-(6-(2,5-dioxo-2,5-dihydro-1H-pyrrol-1-yl)hexanoyl)-3,9-diazaspiro[5.5]undecan-3-yl)-8-oxooctyl)piperazin-1-yl)-2-methylpyrimidin-4-yl)amino)thiazole-5-carboxamide (Compound 2)**

N-(2-chloro-6-methylphenyl)-2-((2-methyl-6-(4-(8-oxo-8-(3,9-diazaspiro[5.5]undecan-3-yl)octyl)piperazin-1-yl)pyrimidin-4-yl)amino)thiazole-5-carboxamide (76 mg, 0.1 mmol, 1 eq) and 6-(2,5-dioxo-2,5-dihydro-1H-pyrrol-1-yl)hexanoic acid (21.12 mg, 0.100 mmol, 1 eq) were dissolved in DMF (1 mL) at room temperature. DIPEA (52.4 microliters, 0.3 mmol, 3 eq) was added, followed by HATU (38.0 mg, 0.1 mmol, 1 eq). After 23 hours, the material was diluted with MeCN (3 mL) and MeOH (3 mL) and purified by preparative HPLC, then lyophilized. The desired product (23.5 mg, 0.023 mmol, 23% yield) was isolated as a white solid.

**$^1$H NMR** (400 MHz, DMSO-d6) δ 11.53 (s, 1H), 9.83 (s, 1H), 8.17 (s, 1H), 7.38 - 7.28 (m, 1H), 7.20 (dd, J = 13.1, 5.5 Hz, 2H), 6.94 (s, 2H), 6.05 (s, 1H), 3.32 (s, 14H), 2.37 (s, 3H), 2.25 - 2.13 (m, 8H), 1.54 (s, 2H), 1.39 (dt, J = 23.5, 7.3 Hz, 12H), 1.32 - 1.19 (m, 11H), 1.19 - 1.07 (m, 3H).

**LCMS**: 915.3 $[M+H]^+$
